# Comparative transcriptome investigation of *Nosema ceranae* infecting eastern honeybee workers

**DOI:** 10.1101/2022.01.12.476029

**Authors:** Yuanchan Fan, Jie Wang, Kejun Yu, Wende Zhang, Zongbin Cai, Minghui Sun, Ying Hu, Xiao Zhao, Cuiling Xiong, Qingsheng Niu, Dafu Chen, Rui Guo

## Abstract

*Apis cerana* is the original host for *Nosema ceranae*, a widespread fungal parasite resulting in bee nosemosis, which leads to severe losses for apiculture industry throughout the world. However, knowledge of *N. ceranae* infecting eastern honeybees is extremely limited. Currently, the mechanism underlying *N. ceranae* infection is still largely unknown. Based on our previously gained high-quality transcriptome datasets, comparative transcriptomic investigation was conducted in this work, with a focus on virulence factor-associated differentially expressed genes (DEGs). Microscopic observation showed that *A. c. cerana* workers’ midguts were effectively infected after inoculation with clean spores of *N. ceranae*. Totally, 1411, 604, and 38 DEGs were identified from NcCK vs. NcT1, NcCK vs. NcT2 and NcT1 vs. NcT2 comparison groups. Venn analysis showed that ten up-regulated genes and nine down-regulated ones were shared by aforementioned comparison groups. GO category indicated these DEGs were involved in a series of functional terms relevant to biological process, cellular component, and molecular function, such as metabolic process, cell part, and catalytic activity. Additionally, KEGG pathway analysis suggested that the DEGs were engaged in an array of pathways of great importance, such as metabolic pathway, glycolysis, and biosynthesis of secondary metabolites. Further, expression clustering analysis demonstrated that majority of genes encoding virulence factors such as ricin B lectins and polar tube proteins displayed apparent up-regulation, whereas a few virulence factor-associated genes such as hexokinase gene and 6-phosphofructokinase gene presented down-regulation during the fungal infection. Finally, the expression trend of 14 DEGs was confirmed by RT-qPCR, validating the reliability of our transcriptome datasets. These results together demonstrated that an overall alteration of the transcriptome of *N. ceranae* occurred during the infection of *A. c. ceranae* workers, and most of virulence factor-related genes were induced to activation to promote the fungal invasion. Our findings not only lay a foundation for clarifying the molecular mechanism underlying *N. ceranae* infection of eastern honeybee workers, but also shed light on developing novel targets for microsporidiosis control.

## 1. Introduction

*Nosema ceranae* is an obligate unicellular fungal parasite that specifically infects bee midgut epithelial cells. *N. ceranae* had been first identified in eastern honeybee (*Apis cerana*) by Fries et al. [1], thereafter it swiftly spread to western honeybee (*Apis mellifera*) colonies reared in Europe and Taiwan province, China [2]. Currently, *N. ceranae* could be detected in colonies all over the world. *N. ceranae* infestation results in a battery negative impact on bee host, such as shortened life span, energy stress, immunosuppression, cell apoptosis inhibition [3,4,5,6], earlier foraging activity, and impaired navigation and cognitive ability [7,8]. A close connection between *N. ceranae* and colony collapse disorder (CCD) had been suggested by several studies [9,10].

*N. ceranae* exists outside the host cell only as dormant spores. After ingestion by the bee host, the inside polar tube is rapidly extruded to pierce the cell membrane followed by injection of the infective sporoplasm via hollow polar tube [1,11,12]. The intracellular life cycle of *N. ceranae* can be divided into two phases including the proliferative phase (merogony) and the sporogonic phase (sporogony), and ends with the formation of spores. Nevertheless, it’s hard to completely isolate *N. ceranae* at above-mentioned two different phases, which is a key factor limiting further study on the fungal parasite during the infection process. Advances in next-generation sequencing technology have allowed a deeper understanding of host response and parasite/pathogen infection. Previously, several studies were conducted to investigate responses of western honeybee workers to microsporidian infestation, Badaoui et al. analyzed the gene expression of *A. m. ligustica* workers at 5, 10, and 15 days post *N. ceranae* infection by RNA-seq technology, and found that the expression of genes encoding host antimicrobial peptides, cuticle proteins, and odor binding proteins were down-regulated, resulting in the decline of immune function of the host, so as to promote the survival and propagation of *N. ceranae* at the colony level [13]; based on fluorescence in situ hybridization (FISH) and immunostaining experiments, Panek et al. investigated the impact of *N. ceranae* on *A. m. ligustica* workers epithelium renewal by following the mitotic index of midgut stem cells during a 22-day *N. ceranae* infection, the results showed that *N. ceranae* can negatively alter the gut epithelium renewal rate and disrupt some signaling pathways involved in the gut homeostasis [14]. Transcriptome analysis of the intestinal tract of *A. c. cerana* exposed to *N. ceranae* demonstrated that microsporidian infection inhibited genes relevant to homeostasis and renewal in wnt signaling pathway [15]. Comparatively, omics study on the *N. ceranae* infecting bee hosts is very limited, and the underlying mechanism of *N. ceranae* infection is still vague.

In our previous work, we performed RNA sequencing of clean spores of *N. ceranae*, followed by transcriptome-wide identification of ncRNAs such as miRNAs, lncRNAs, and circRNAs in *N. ceranae* spores [16, 17, 18]. Recently, we conducted deep sequencing of the midguts of *A. c. cerana* workers at 7 and 10 days post inoculation (dpi) with *N. ceranae* spores, and deciphered the host cellular and humoral responses to fungal infection [19]; in addition, we analyzed the expression profile of highly expressed genes (HEGs) and discussed their potential roles in *N. ceranae* infestation [20]. In view of that transcriptomic study on *N. ceranae* during the infection process is still very lagging, in the present study, based on the obtained transcriptome datasets, we first filtered out the parasite-derived data and then performed comparative investigation combined with transcriptome data derived from *N. ceranae* spores, followed by deep investigation of the dynamics of genes in *N. ceranae* as well as virulence factor-associated pathways and genes. To the best of our knowledge, this is the first report of omics study on *N. ceranae* invading eastern honeybee.

## 2. Results

### 2.1. Verification of infection of *A. c. cerana* worker by *N. ceranae*

Under optical microscope, oval and highly refractive dispersed spores were observed **(Figure 1A)**. Further, AGE indicated that expected fragment (approximately 76 bp) was amplified from the purified spores with specific primers for *N. ceranae*, while no signal band was detected using specific primers for *N. apis* **(Figure 1B)**. These results verified that the purified spores were indeed *N. ceranae* spores.

**Figure 1.**
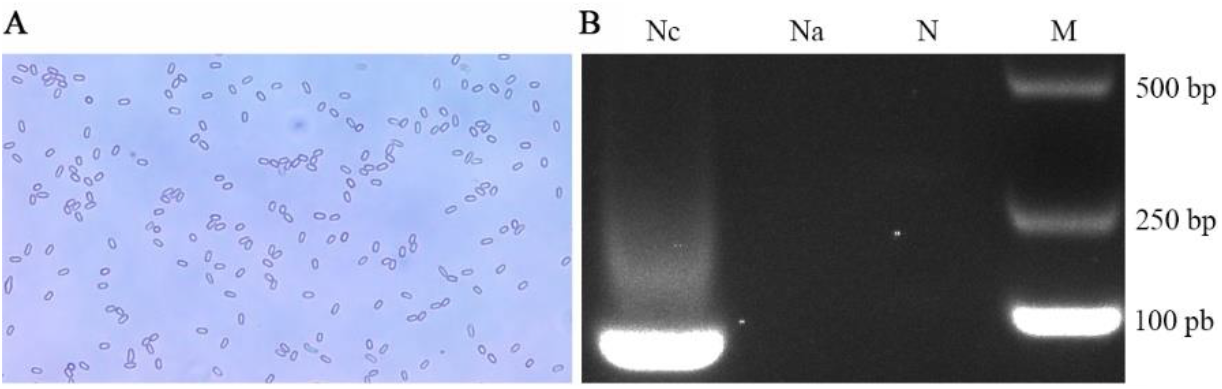
Microscopic detection and PCR validation of *N. ceranae* spores. (A) Microscopic detection (400 times amplification). (B) AGE for PCR amplified fragments, Lane Nc: Specific primers for *N. ceranae*, Lane Na: Specific primers for *N. apis*, Lane N: Sterile water (Negative control), Lane M: DNA marker.

Based on microscopic observation of paraffin sections, it’s found that there were a number of *N. ceranae* spores in the *A. c. cerana* worker’s midgut epithelial cells at 11 dpi with *N. ceranae* **(Figure 2A-B)**, whereas no fungal spores can be detected in the worker’s midgut epithelial cells at 11 dpi without *N. ceranae* **(Figure 2C-D)**. In addition, the structure of midgut epithelial cells of *N. ceranae*-infected worker was fragmentary and the nucleic acid substances were dispersed and unclear **(Figure 2A-B)**, while that of un-infected worker was intact and the deeply colored cell nucleus were visible **(Figure 2C-D)**. The results together suggested that the *A. c. cerana* workers were infected by *N. ceranae* and the host midgut epithelial cell structure was destructed by the fungal infection.

**Figure 2.**
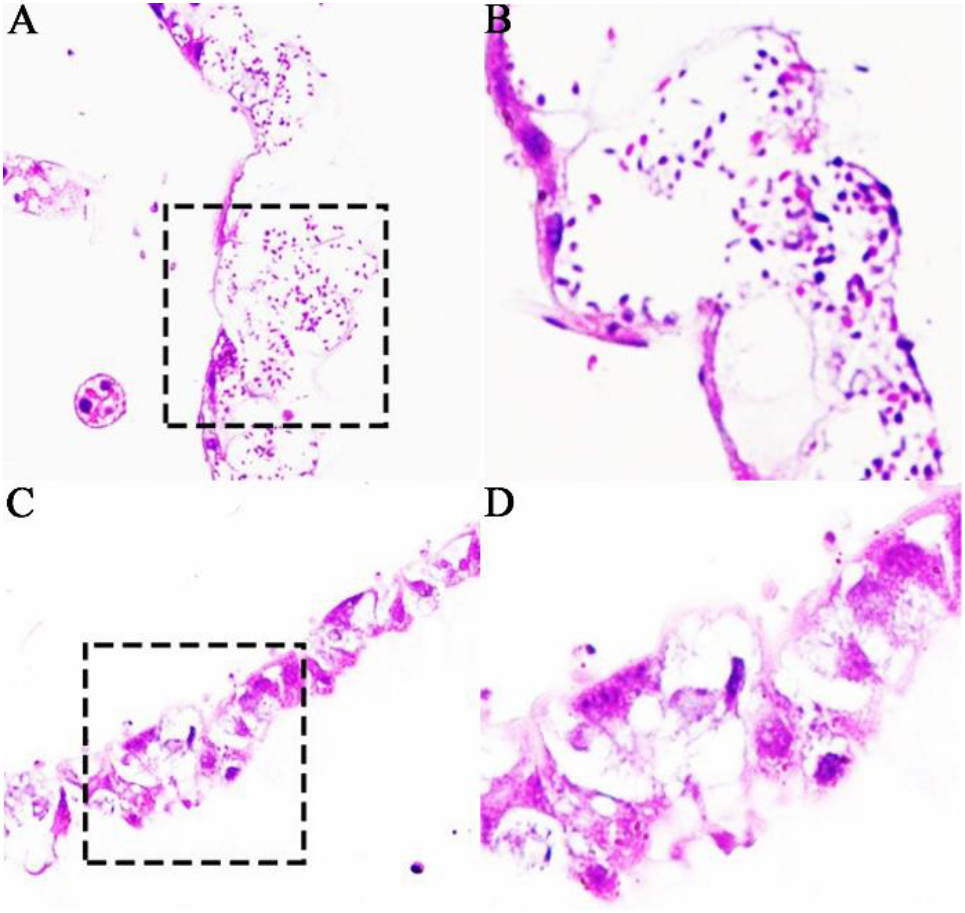
Microscopic observation of paraffin sections of *N. ceranae*-inoculated and un-inoculated *A. c. cerana* workers’ midguts. (A) Worker’s midgut at 11 dpi without *N. ceranae* under 200 times amplification; (B) Worker’s midgut at 11 dpi without *N. ceranae* under 400 times amplification. Black dashed box shows the region for observation under 400 times amplification.

### 2.2. Differential gene expression profile of *N. ceranae* infecting *A. c. cerana* workers

Totally, 1411, 604, and 38 DEGs were identified in NcCK *vs*. NcT1, NcCK *vs*. NcT2, and NcT1 *vs*. NcT2 comparison groups, respectively. The numbers of up-regulated genes were 711, 240, and 17, while those of down-regulated genes were 700, 360, and 21, respectively **(Figure 3A)**. Additionally, Venn analysis showed that there were 10 and nine shared up- and down-regulated genes in the aforementioned three comparison groups, whereas the numbers of unique up-regulated (down-regulated) genes were 417 (354), two (20), and five (10), respectively **(Figure 3B)**.

**Figure 3.**
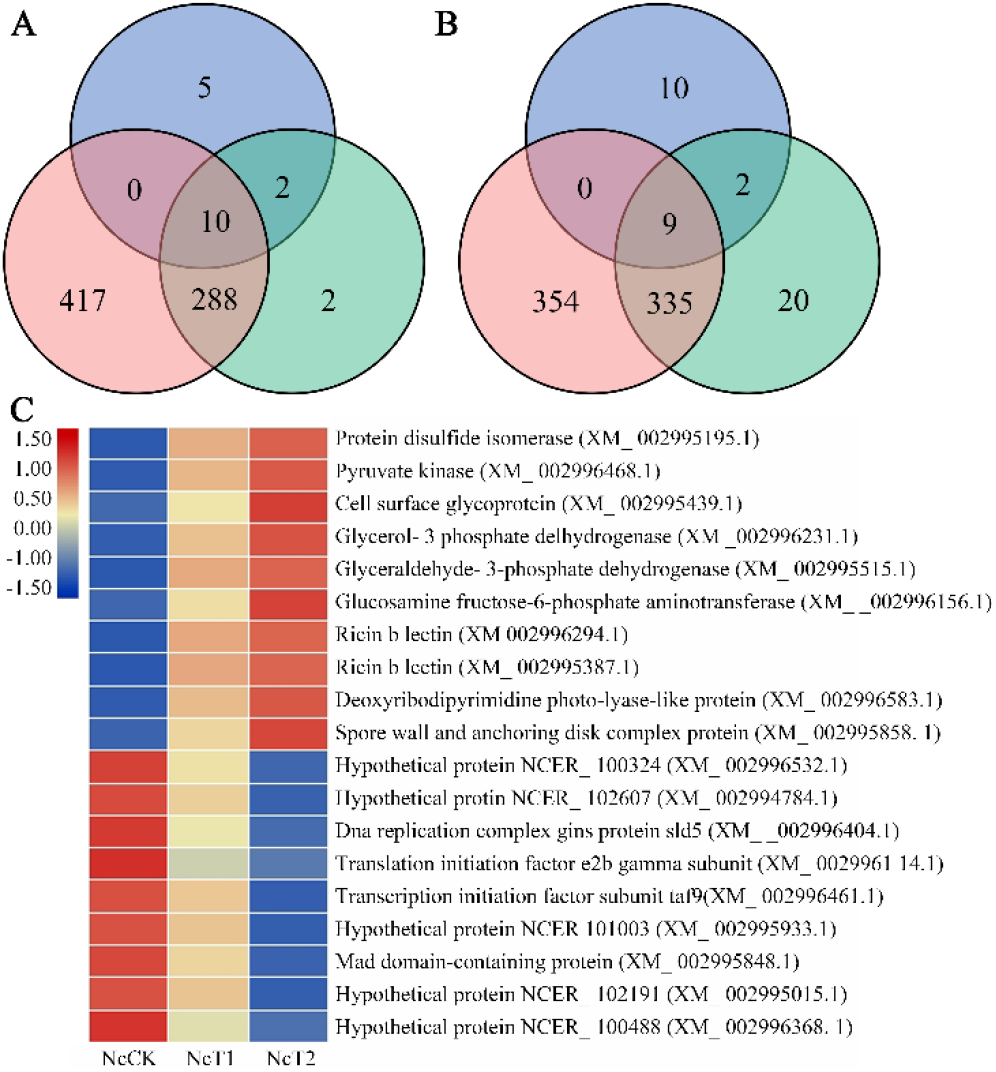
Analysis of DEGs in *N. ceranae*. (A) Venn analysis of up-regulated genes in NcCK *vs*. NcT1, NcCK *vs*. NcT2, and NcT1 *vs*. NcT2 comparison groups. (B) Venn analysis of down-regulated genes in NcCK *vs*. NcT1, NcCK *vs*. NcT2, and NcT1 *vs*. NcT2 comparison groups. (C) Heatmap of common DEGs in NcCK, NcT1, and NcT2 groups.

### 2.3. Function and pathway annotation of DEGs in *N. ceranae* infesting *A. c. cerana* workers

GO term analysis suggested that the DEGs in NcCK *vs*. NcT1 comparison group were engaged in 638 biological process-associated functional terms such as metabolic process, single-organism process, and cellular process; 377 cellular component-associated terms such as cell, cell part, and organelle; and 424 molecular function-associated terms such as catalytic activity, transport, and binding **(Figure 4A)**. In NcCK *vs*. NcT2 comparison group, the DEGs could annotate to 22 GO terms, including 291 biological process-related items such as metabolic process and single-organism, 160 cellular component-related terms such as cell and cell part, and 237 molecular function-related items such as catalytic activity and nucleic acid binding transcription factor activity **(Figure 4B)**. The DEGs in NcT1 *vs*. NcT2 comparison group were involved in seven functional terms such as metabolic process, cell, and catalytic activity **(Figure 4C)**.

**Figure 4.**
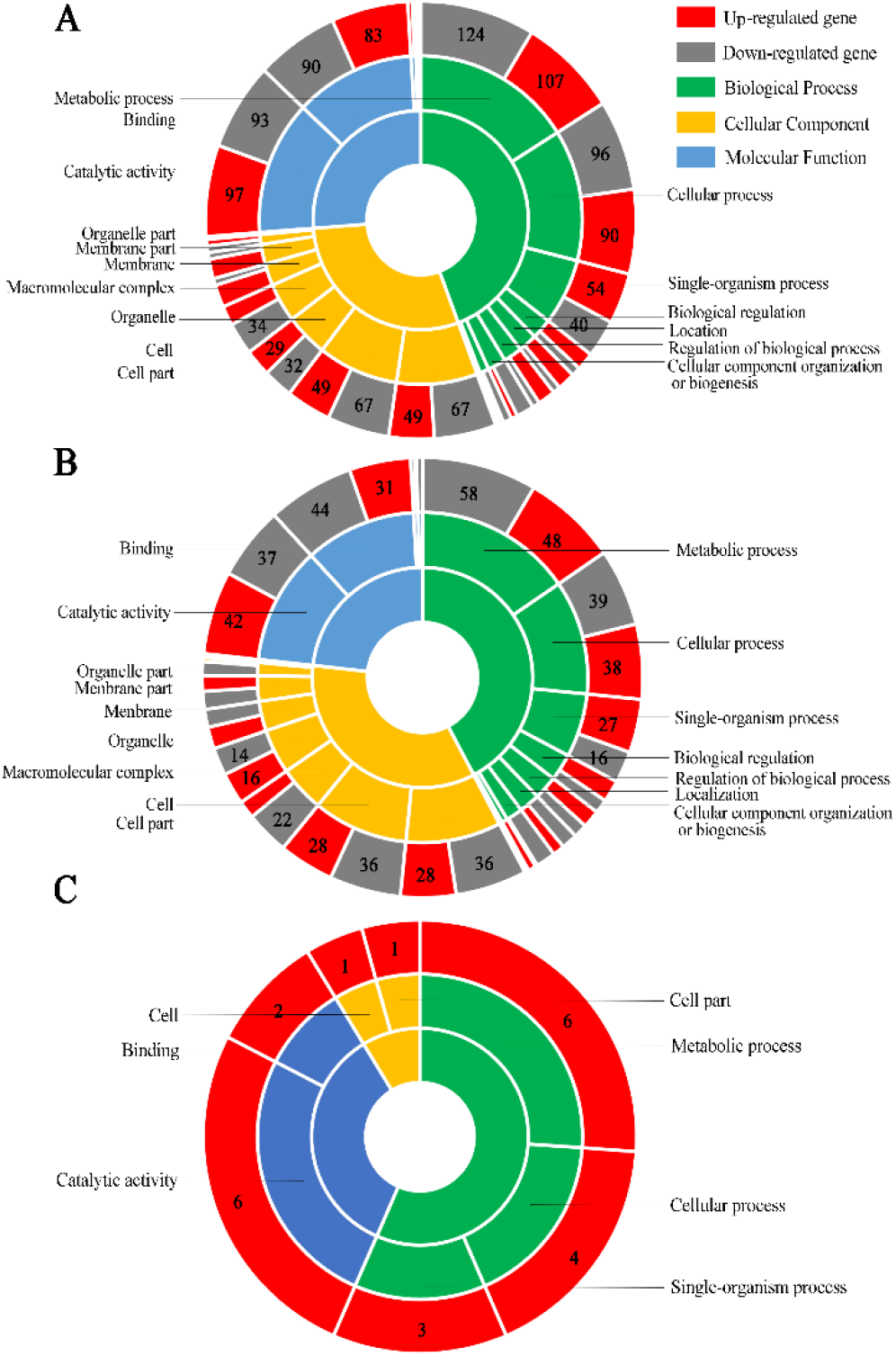
GO category of DEGs. (A) DEGs in NcCK *vs*. NcT1. (B) DEGs in NcCK *vs*. NcT2; (C) DEGs in NcT1 *vs*. NcT2.

In addition, KEGG pathway analysis indicated that the DEGs in NcCK *vs*. NcT1 comparison group were relevant to 241 pathways, among these the most abundant groups were metabolic pathway, biosynthesis of secondary metabolites, ribosome, ribosome biogenesis in eukaryotes, and biosynthesis of antibiotics **(Figure 5A)**. In NcCK *vs*. NcT2 comparison group, the DEGs were relative to 201 pathways including metabolic pathway, biosynthesis of secondary metabolites, ribosome, ribosome biogenesis in eukaryotes, and biosynthesis of antibiotics **(Figure 5B)**. The DEGs in NcT1 *vs*. NcT2 comparison group were associated with 32 pathways such as metabolic pathway, biosynthesis of antibiotics, biosynthesis of secondary metabolites, glycolysis/gluconeogenesis, and microbial metabolism in diverse environments **(Figure 5C)**.

**Figure 5.**
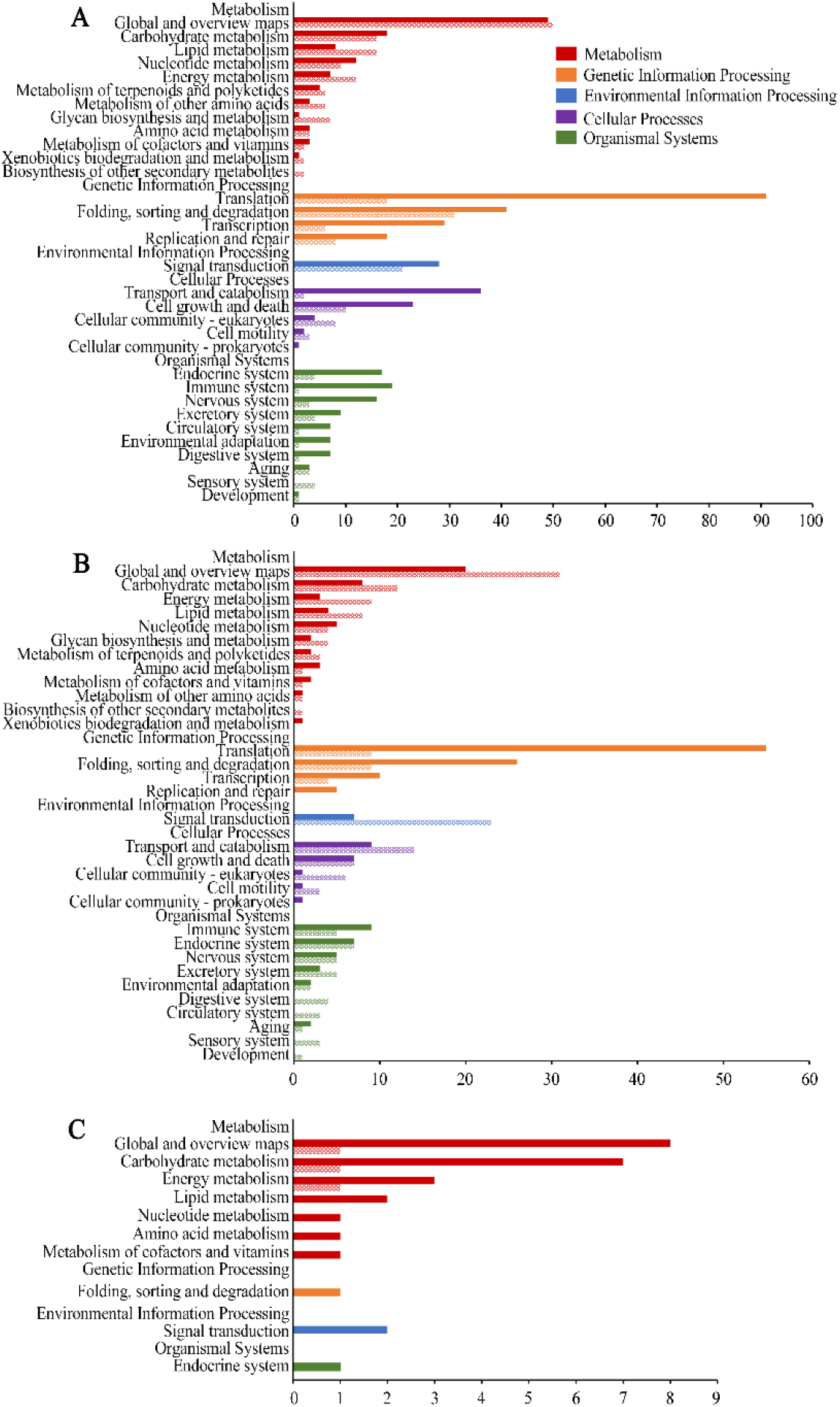
Pathways annotated by DEGs. (A) DEGs in NcCK *vs*. NcT1. (B) DEGs in NcCK *vs*. NcT2. (C): DEGs in NcT1 *vs*. NcT2.

### 2.4. Virulence factor-associated DEGs in *N. ceranae* invading *A. c. cerana* workers

Further investigation was conducted to explore virulence factor-associated DEGs in above-mentioned comparison groups, a total of 20 DEGs were identified, including six spore wall protein coding genes, three ricin B lectin protein coding genes, three ATP/ADP translocase protein coding genes, two polar tube protein coding genes, two ABC transporter protein coding genes, one chitin synthase protein coding gene, one 6-phosphofructokinase protein coding genes, one hexokinase protein coding gene, and one pyruvate kinase protein coding gene. Moreover, the expression clustering showed that majority of virulence factor-encoding genes were induced to activation during the infection process, whereas a few were suppressed to a large externt **(Figure 6)**.

**Figure 6.**
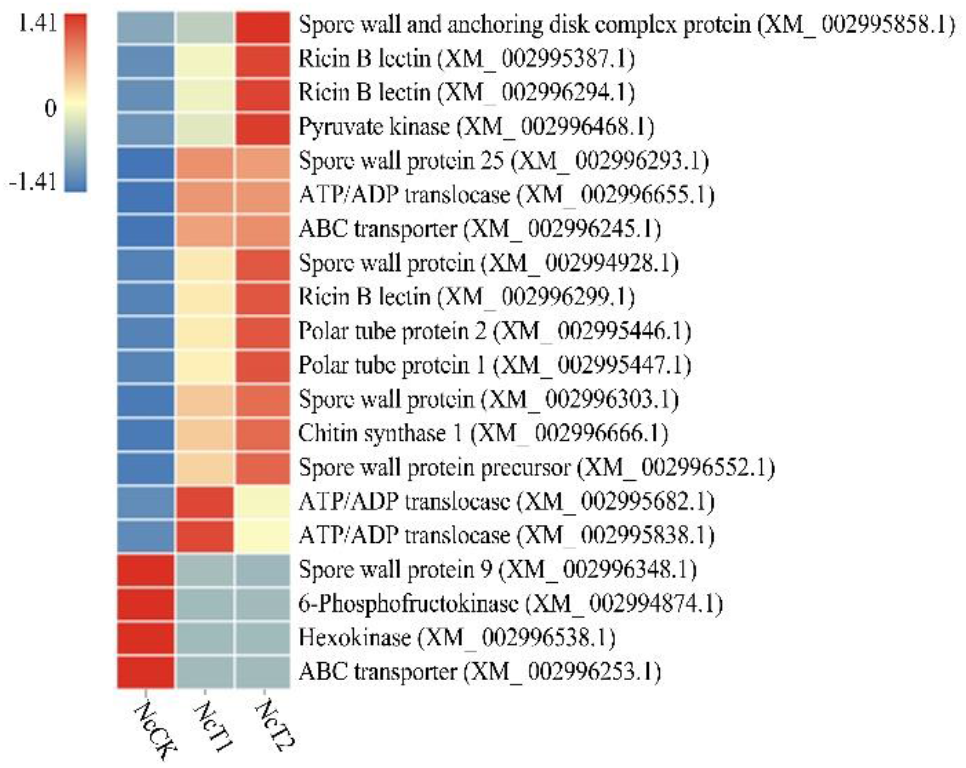
Heatmap of virulence factor-associated DEGs shared by NcCK, NcT1, and NcT2 groups.

### 2.5. Verification of DEGs via RT-qPCR

Fifteen DEGs were randomly selected for RT-qPCR validation, the result suggested that the expression trend of 14 were consistent with those in transcriptome data **(Figure 7A-B)**, confirming the reliability of the sequencing data used in this current work.

**Figure 7.**
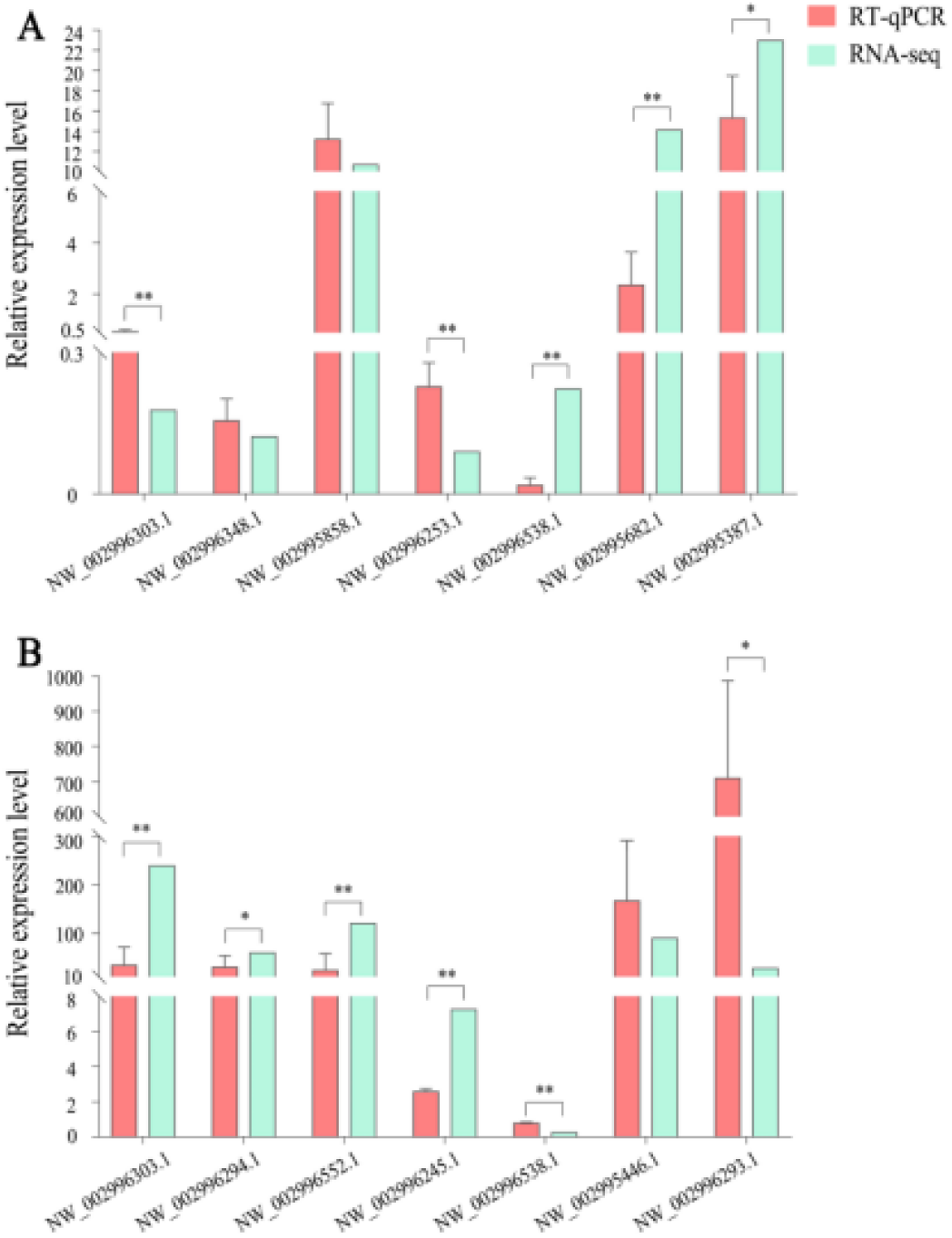
RT-qPCR verification of DEGs in *N. ceranae* infecting *A. c. cerana* workers. (A) DEGs in NcCK *vs*. NcT1. (B) DEGs in NcCK *vs*. NcT2. Error bars represent the variance of RT-qPCR results of each DEG. Bars with asterisk symbol indicate statistical differences (P < 0.05).

## 3. Discussion

Currently, the mechanism regulating *N. ceranae* infestation remains to be clarified. Here, based on our previously obtained transcriptome data from *N. ceranae*-inoculated and un-inoculated *A. c. cerana* workers’ midguts at 7 dpi and 10 dpi, the mixed transcriptome data were first filtered to gain those *N. ceranae*-derived data, and then used for comparative analysis in conjunction with transcriptome data from *N. ceranae* spores. Following the similar technique protocol, our them previously investigated transcriptomic dynamics of *N. ceranae* invading *A. m. ligustica* workers and *Ascosphaera apis* infecting *A. m. ligustica* or *A. c. cerana* larvae [30,31,33-35], and the protocol was verified to be feasible and reliable, offering a reference for investigating pathogens or endoparasites during the infection process, especially those hardly isolated from hosts under current technical conditions.

In NcCK *vs*. NcT1 and NcCK *vs*. NcT2 comparison groups, 1411 and 604 DEGs were respectively identified, including 657 and 240 up-regulated genes as well as 700 and 364 down-regulated ones. This indictive of the overall alteration of genes in *N. ceranae* during the infection process. GO term analysis showed that the aforementioned DEGs were involved in 24 and 22 functional terms associated with biological processes such as metabolic process and cellular process, cellular components such as cell and organelle, and molecular function such as catalytic activity and transport. In addition, these DEGs were annotated to 241 and 201 pathways, respectively, including material metabolism-related pathways such as Corban metabolism and energy mechanism-related pathways such as glycolysis/gluconeogenesis. The response of fungal cells to external signals is regulated by mitogen-activated protein kinase (MAPK), which phosphorylates many downstream proteins and further alter gene expression, followed by participation in various process such as proliferation, differentiation, and apoptosis [36]. In this current work, three and two DEGs in NcCK *vs*. NcT1 and NcCK *vs*. NcT2 were found to enrich in MAPK signaling pathway, among which XM_002995908.1 (log2FC= 4.44, 4.93) and XM_002996683.1 (log2FC= 4.63, 4.51) were shared by the both comparison groups, while there was a unique DEG (XM_002996282.1, log2FC=-1.01) in NcCK *vs*. NcT1. This indicated that MAPK signaling pathway was significantly activated during *N. ceranae* infection and used by the fungal parasite to response to the internal environment of epithelial cells in *A. c. cerana* workers’ midguts. Previously, we conducted transcriptomic investigation of *A. m. ligustica* workers’ midguts responding to *N. ceranae* invasion, and detected that seven (GenBank accession no. 9423510, 424610, 9424462, 9422987, 9422306, 9411598 and 9424007) DEGs in midguts at 7 dpi and 10 dpi were enriched in MAPK signaling pathway [31], which suggested that the MAPK pathway regulated *A. m. ligustica* immune response to *N. ceranae*. These results together demonstrated that MAPK signaling pathway was likely to play a vital part in *N. ceranae* infection, but there were differences of this signaling pathway in *N. ceranae* when infecting different bee species.

Ricin, a kind of toxic heterodimer protein in castor seed, has a ricin chain B (RTB) that connects a ricin chain A (RTA) via a disulfide bond [37]. Lectins are proteins that recognize glycan ligand. As virulence factors of many pathogens, lectins exert function by mediating cell adhesion, innate immune defense, and pathogen infection [38]. The binding of ricin B lectin (RBL) contributes to microbial infection and promotes pathogen attachment or entry into host cells. NBRBL3, a lectin protein of *Nosema bombyx*, was confirmed to be secreted from the spore during fungal proliferation and then mediate host cell recognition and adhesion [38]. In this research, three ricin B lectin-associated DEGs were observed in both NcCK vs. NcT1 and NcCK *vs*. NcT2 comparison groups, including XM_002995387.1 (log2FC=4.52, 5.67), XM_002996294.1 (log2FC= 4.66, 5.88), and XM_002996299.1(log2FC =2.31, 2.96), and all of these were apparently up-regulated, implying that *N. ceranae* may synthesize and secrete ricin B lectin to increase the adhesion between spores and host cells to promote fungal proliferation [31].

There is a hard spore wall outside the microsporidia, which is composed of high electron density protein outer spore wall layer, low electron density chitin inner spore wall layer, and fibrous plasma membrane. Spore wall protein (SWP) is proved to be a crucial virulence factor of microsporidia, which can recognize and adhere to host cells during microsporidian infection [39]. The spore wall protein SWP25 of *Nosema bombyx* was located in the endospore layer with a signal peptide, which was suggested to be implicated in chitin interaction of endospore powder and the construction of spore wall via the heparin binding motif (HBM) [40]. Here, one SWP25-encoding gene (XM_002996293.1) showed an up-regulation trend in host midguts at 7 dpi (log2FC=4.82) and 10 dpi (log2FC=4.77), suggestive of the participation of SWP25 in *N. ceranae* infection of *A. c. cerana* workers. Yang et al. analyzed the interaction between *Nosema bombycis* SWP9 (NbSWP9) with polar tube proteins PTP1 and PTP2 by molecular methods such as co-immunoprecipitation and yeast two-hybrid assay, and found that NbSWP9 was mainly distributed in the polar tubes [41]. In this work, one gene encoding SWP9 (XM_002996348.1) was down-regulated in workers’ midguts at both 7 dpi (log2FC=-3.04) and 10 dpi (log2FC=-3.40), indicating that *A. c. cerana* workers may inhibit the interaction between SWP9 with PTPs via host-parasite interaction. However, additional work is required to decipher the underlying mechanism. In addition, another two SWP encoding genes (XM_002996303.1, log2FC=7.45, 7.90; XM_002994928.1, log2FC =5.08, 5.82), one SWP precursor encoding gene (XM_002996552.1, log2FC =6.36, 6.90), and one spore wall and anchoring disk complex protein encoding gene (XM_002995858.1, log2FC =3.41, 5.80) were identified and all of them were observed to up-regulate in host midguts at both 7 dpi and 10 dpi, which demonstrated that these SWP-associated genes were activated to increase the synthesis of corresponding SWPs, further enhancing the N. ceranae invasion.

For microsporidia, the infective protoplasm is transferred from the spore to host cell through the polar tube, further starting the proliferation process. Therefore, the polar tube protein is also considered as a virulence factor of importance. Yang et al. discovered that the polar tube and spore wall of microsporidia are the main components of mature spores adhering to and infecting the host cells [42]. Long et al. performed lectin blotting and β-elimination reaction of NbPTP1 in *N. bombycis*, the results showed that NbPTP1 had O-glycosylation modification characteristics, which was conducive to adhesion, infection, and maintenance of the stability of the polar tube. Moreover, the PTP1 protein sequence contains high proline, which can improve the elasticity of the polar tube [43]. Here, two PTP1 encoding genes shared by the aforementioned two comparison groups were found to be up-regulated, including XM_002995447.1 (log2FC = 6.01, 6.84) and XM_002995446.1 (log2FC =5.71, 6.48), indicating that the tubulin was engaged in the infection process of *N. ceranae* and played a role in the maintenance of the stability of polar tube.

Chitin is a major component of the spore wall of *N. ceranae*. Chitin synthase is an important enzyme implicated in chitin biosynthesis. Cabib et al. discovered that chitin synthase 1 and chitin synthase 2 exerted synergistic function in chitin repair in yeast, and jointly participated in the cell wall construction [44]. In the present study, the chitin synthase 1 encoding gene (XM_002996666.1) was up-regulated in both NcCK *vs*. NcT1 (log2FC = 4.07) and NcCK *vs*. NcT2 (log2FC = 4.52) comparison groups, implying that it played a part in the construction and repair of spore wall to assist the *N. ceranae* infection. However, we did not detect the differential expression of chitin synthase gene in *N. ceranae* infecting the *A. m. ligustica* workers. It’s speculated that chitin synthesis gene was a special virulence factor for N. ceranae, playing different roles when infecting different bee species.

Owing to the specific life cycle, microsporidia lost the typical mitochondria during the long-term evolution, which was replaced by mitosis, an organelle that produces a small amount of ATP via glycolysis to meet the basic physiological needs [11]. Accordingly, microsporidia highly depend on stealing energy from the host cells, and glycolysis is the main manner for *N. ceranae* to produce ATP at a pretty low efficiency [12]. Huang et al. conducted transcriptomic analysis of the *N. ceranae* during the 6 days infection cycle, and detected that the genes encoding three rate-limiting enzymes involving catalyzing glycolysis pathway continued to up-regulate, including hexokinase, pyruvate kinase, and phosphofructokinase; and at the end of infection (6 dpi), the gene expression levels of ATP/ADP translocase gene, ABC transporter gene, and catalytic glycolysis encoding gene were plunged [45]. Hexokinase catalyzes glucose phosphorylation to form glucose-6-phosphate (g-6-p), and the expression of hexokinase is regulated by ADP. Pyruvate kinase can catalyze the conversion of phosphoenolpyruvate to pyruvate and further enhance the production of ATP. Fructose-6-phosphate (f-6-p) is hydrolyzed by 6-phosphofructokinase, glucose-1 (F-1) and 6-diphosphate (6-2p) were inhibited by ATP with high concentration [56-48]. Noticeably, the pyruvate kinase encoding gene XM_002996468.1 (log2FC = 2.81, 3.92) and hexokinase encoding gene XM_002995838.1 (log2FC = 3.78, 3.23) were up-regulated in NcCK *vs*. NcT1 and NcCK *vs*. NcT2, which suggested that activation of pyruvate kinase and hexokinase was a strategy of *N. ceranae* during the infection of *A. c cerana* workers. However, the 6-phosphofructokinase encoding gene XM_002994874.1 (log2FC =-4.72, - 4.03) was down-regulated, which was inconsistent with previous study [45]. We inferred that it might be suppressed by ATP with high concentration. The underlying mechanism needs to be further explored.

ABC transporters represent the largest family of transmembrane proteinsm which contribute to transportation of ATP. Most of ABC transporters rely on ATP binding and hydrolysis to transport amino acids, lipids, sugars, peptides, ions, and other substrates from the cell fluid to intracellular or extracellular region [49]. He et al. identified 234 ABC transporters from 18 microsporidium genomes and divided them into five subfamilies: ABCBs, ABCCs, ABCEs, ABCFs, and ABCGs. Two subfamilies of ABCBs and ABCDs were lost in microsporidian genome. On the basis of qPCR, Western blotting, and RNAi, He et al. revealed that NoboABCG1.1 regulated the fungal reproduction and promoted the proliferation of *N. bombycis* [50]. Here, the expression trend of two ABC transporter-encoding genes was altered in the above-mentioned two comparison groups; XM_002996245.1 (log2FC =2.52, 2.86) was up-regulated, while XM_002996253.1 (log2FC =-3.49, −3.68) was down-regulated. This indicated that ABC transporters may play an essential role in the transmembrane transport of materials and energy during the *N. ceranae* infection, and different ABC transporters encoding genes may have different roles in this process.

ATP/ADP translocase transports ATP synthesized in mitochondrial matrix to cytoplasm for cell utilization in eukaryotes. *N. ceranae* needs to steal ATP from host cells by ATP/ADP translocase to meet its growth and development needs. Paldi et al. found four ATP/ADP translocases in *N. ceranae* infecting *A. mellifera* workers based on genomic sequencing and investigation, and concluded that ATP/ADP translocases may play a role in transporting ATP from hose cells to the mitosis in *N. ceranae* [51]. In the present study, the up-regulation of two ATP/ADP translocase genes was detected in NcCK *vs*. NcT1 and NcCK *vs*. NcT2, including XM_002995682.1(log2FC =3.82, 3.33), XM_002996655.1(log2FC =2.54, 2.54); whereas another gene XM_002996538.1 (log2FC =-2.08, −2.08) was down-regulated, indicative of the complex interaction between microsporidian and host. The result supported the vital function of ATP/ADP transporters in energy acquisition of *N. ceranae* invading *A. c. cerana* workers.

## 4. Materials and Methods

### 4.1. Fungal spore and honeybee

Clean spores of N. ceranae were previously purified using Percoll discontinuous gradient centrifugation protocol and preserved in Honeybee Protection Laboratory [18], College of Animal Sciences (College of Bee Science). A. c. cerana workers were selected from three colonies located in the teaching apiary of the College of Animal Sciences (College of Bee Science) in Fujian Agriculture and Forestry University. No Varroa mite was observed during the whole study. The disappearance of seven common bee viruses (KBV, IAPV, ABPV, DWV, SBV, BQCV, and CBPV) and two bee microsporidia (Nosema apis and N. ceranae) in the newly emergent A. c. cerana workers was verified with RT-PCR assay [21].

### 4.2 Microscopic observation and PCR validation of *N. ceranae spores*

The prepared spores of N. ceranae was subjected to microscopic observation using an optical microscope (SIOM, Shanghai, China). Further, total DNA of spores were isolated and used as templates for reverse transcription; the resulting cDNA was then used as templates for PCR amplification with previously described specific primers for N. ceranae and N. apis [22-23]; the amplified products were detected by 1.5% agarose gel electrophoresis (AGE). Sterile water was set as a negative control.

### 4.3 Preparation and detection of paraffin section of honeybee midgut tissue

One-day-old workers of A. c. cerana in N. ceranae-inoculated groups were each artificially fed with 5 ul 50% (w/v) sucrose solution containing 1 × 106 spores; while 1 -old-day workers in un-inoculated groups were each fed with 5 μL of 50% (w/v) sucrose solution without spores. At 11 dpi with N. ceranae spores, the midgut tissues in the N. ceranae-inoculated and un-inoculated groups were respectively harvested and fixed with 4% paraformaldehyde. According to our previously described protocol [21], on the basis of a microtome (Leica, Nussloch, Germany) and an embedding center (Junjie, Wuhan, China), paraffin sections were prepared and then stained with hematoxylin eosin (HE) stain by Shanghai Sangon Biological Engineering Co. Ltd, followed by detection utilizing an optical microscope with digital camera (SOPTOP, Shanghai, China).

### 4.4 Transcriptome data source

Midgut tissues of A. c. cerana workers at 7 dpi and 10 dpi with *N. ceranae* spores and corresponding un-inoculated worker’s midgut tissues were previously prepared and sequenced on Illumina HiSeqTM 4000 platform with a stand-specific cDNA library-based strategy [17]. The raw datasets had been deposited in the National Biotechnology Information Center (NCBI) SRA database (https://www.ncbi.nlm.nih.gov/sra) under BioProject number: PRJNA395264. Quality control of the produced raw reads were previously conducted according to the method described by Chen et al [17,24]. Briefly, raw reads containing adapters, more than 10% of unknown nucleotides (N), and more than 50% of low quality (q value ≤ 20) bases were removed to gain high-quality clean reads, which were then mapped to the ribosome RNA (rRNA) database (www.arb-silva.de/) (20 March 2021) using Bowtie2 software [24]. The result indicated that 174, 700, 032, 205, 297, 946, 124, 216, 829, and 99, 030, 788 raw reads were generated, and after quality control, 171, 868, 061, 200, 570, 776, 121, 949, 977, and 97, 432, 267 clean reads were gained, with Q20 of 94.96%, 94.58%, 94.60%, and 94.91% respectively [25]. Hence, the high-quality RNA-seq data can be used for transcriptomic analyses in the present study. In another work, we conducted deep sequencing of clean spores of N. ceranae (NcCK) using Illumina HiSeq-based RNA-seq [18]. A total of 416, 156, 600 raw reads were produced, and after quality control, 210, 824, 312 clean reads were obtained, with average Q30 of 92.63%, which suggested that the high-quality transcriptome data can be used in this study. The relevant raw data had been deposited in NCBI SRA database (https://www.ncbi.nlm.nih.gov/sra) under BioProject number: PRJNA562784.\

### 4.5 Identification and analysis of DEGs in *N. ceranae*

Following the standard of P value ≤ 0.05 and | log2(Fold change) | ≥ 1, the DEGs in NcCK *vs*. NcT1 and NcCK *vs*. NcT2 comparison groups were screened by edgeR software [26]. Venn analysis of up- and down-regulated genes in these comparison groups and expression cluster analysis were carried out based on OmicShare platform (https://www.omicshare.com/). GO (Gene Ontology) categorization of DEGs were carried out using WEGO software [27]. Blastall tool was employed to conduct pathway analysis by comparing DEGs against KEGG (Kyoto Encyclopedia of Genes and Genomes) database (https://www.kegg.jp/) [28].

### 4.6 Investigation of virulence factor-associated DEGs

On the basis of associated documentations with N. ceranae and findings from our previous studies on N. ceranae [16-20, 23, 29-31], virulence factors such as spore wall protein, ricin B lectin, chitinase, polar tube protein, phosphofructokinase, ATP/ADP translocase, ABC transporter, hexokinase, and pyruvate kinase and associated DEGs were selected for further investigation. Expression clustering analysis of aforementioned DEGs was performed utilizing OmicShare platform.

### 4.7 RT-qPCR validation of DEGs

Fifteen DEGs were randomly selected from NcCK *vs*. NcT1 (XM_002994928.1, XM_002996348.1, XM_002996299.1, XM_002996655.1, XM_002996468.1 and XM_002996253.1), NcCK vs. NcT2 (XM_002996303.1, XM_002996294.1, XM_002995682.1, XM_002996538.1, XM_002996655.1 and XM_002996253.1), and NcT1 *vs*. NcT2 (XM_002996294.1 and XM_002996468.1) comparison groups and subjected to RT-qPCR validation. actin gene (gene6001) was used as an internal reference. Specific forward and reverse primers for these DEGs and actin were designed with primer premier 5 (Table SI). Total RNA of *N. ceranae* spores and *N. ceranae*-inoculated workers’ midguts at 7 dpi and 10 dpi were respectively isolated using RNA Extraction Kit (TaKaRa company, Dalian, China). cDNA was synthesized through reverse transcription with oligo dT primer and used as templates for qPCR assay, which was carried out on a QuanStudio RealTime PCR System (ThemoFisher, Walthem, MA, USA). The qPCR reaction was conducted according to the instructions of SYBR Green Dye Kit (Vazyme company, Shanghai, China). Cycling parameters were as follows: 95 °C for 1 min, followed by 40 cycles at 95 °C for 15 s, 55 °C for 30 s, and 72 °C for 45 s. The relative gene expression was calculated based on 2–ΔΔCt method [32]. The experiment was performed three times utilizing three independent biological samples.

### 2.8. Statistical analysis

Statistical analyses were conducted with SPSS software (IBM, Amunque, NY, USA) and GraphPad Prism 7.0 software (GraphPad, San Diego, California, USA). Data were presented as mean ± standard deviation (SD). Statistics analysis was performed using Student’s t-test and one-way ANOVA. Additionally, significant *(p < 0*.*05)* GO terms and KEGG pathways were filtered by performing Fisher’s exact test with R software 3.3.1.

## 5. Conclusions

In summary, our results demonstrated that the overall transcriptome dynamics of N. ceranae was altered during the infection of A. c. cerana workers, a number of virulence factor-associated genes were induced to activation to facilitate the fungal infestation, but some other genes encoding virulence factors were suppressed via host-microsporidian interaction. Findings in this current work not only provide a basis for clarifying the mechanism underlying the N. ceranae infection of A. c cerana workers, but also shed light on the development of novel strategy for nosemosis control.

## Supplementary Materials

Table S1 Primers for RT-qPCR validation performed in this study.

## Author Contributions

DFC and RG designed this study. RG, YCF, JW, KJY, WDZ, ZBC, MHS, YH, XZ and CLX performed bioinformatic analysis and molecular experiment. DFC, QSN and RG supervised the work and contributed to preparation of the manuscript.

## Acknowledgments

We thank all editors and reviewers for their helpful and constructive comments.

